# β-catenin inhibition disrupts the homeostasis of osteogenic/adipogenic differentiation leading to the development of glucocorticoid-induced osteonecrosis of femoral head

**DOI:** 10.1101/2023.10.04.560853

**Authors:** Chenjie Xia, Huihui Xu, Liang Fang, Jiali Chen, Wenhua Yuan, Danqing Fu, Xucheng Wang, Bangjian He, Luwei Xiao, Chengliang Wu, Peijian Tong, Di Chen, Pinger Wang, Hongting Jin

## Abstract

Glucocorticoid-induced osteonecrosis of the femoral head (GONFH) is a common refractory joint disease characterized by bone damage and the collapse of femoral head structure. However, the exact pathological mechanisms of GONFH remain unknown. Here, we observed abnormal osteogenesis and adipogenesis associated with decreased β-catenin in the necrotic femoral head of GONFH patients. In vivo and in vitro studies further revealed that glucocorticoid exposure disrupted osteogenic/adipogenic differentiation of bone marrow mesenchymal cells (BMSCs) by inhibiting β-catenin signaling in glucocorticoid-induced GONFH rats. Col2^+^ lineage largely contributes to BMSCs, and was found an osteogenic commitment in the femoral head through 9 months of lineage trace. Specific deletion of *β-catenin* in Col2^+^ cells shifted their commitment from osteoblasts to adipocytes, leading to a full spectrum of disease phenotype of GONFH in adult mice. Overall, we uncover that β-catenin inhibition disrupting the homeostasis of osteogenic/adipogenic differentiation contributes to the development of GONFH, and identify an ideal genetic modified mouse model of GONFH.

## Introduction

Glucocorticoids, potent immunity regulators, are widely used in the treatment of various autoimmune diseases such as systemic lupus erythematosus (SLE), nephrotic syndrome (NS) and rheumatoid arthritis (RA)^[1]^. However, excessive usage of glucocorticoids has been reported to cause severe adverse effects, especially femoral head osteonecrosis in the joint^[2, 3]^. As a destructive joint disease, glucocorticoid-induced osteonecrosis of the femoral head (GONFH) disables about 100,000 Chinese and more than 20,000 Americans annually^[4, 5]^, bringing a huge financial burden on the society. Currently, there is no ideal medical treatment for GONFH due to its unclear pathological mechanisms. Most patients eventually have to undergo a joint replacement surgery^[6]^. Therefore, in-depth study of GONFH pathogenesis is urgently needed.

Previous *ex-vivo* studies on human necrotic femoral heads revealed prominent pathological features of GONFH including fat droplet clusters, trabecular bone loss, empty lacunae of osteocyte at the early stage and extra subchondral bone destruction, structure collapse at the late stage^[7, 8]^. Glucocorticoid-induced animal models are commonly used to study GONFH pathogenesis, which mimic the early necrotic changes of GONFH but without structure collapse^[9]^. Several hypotheses, such as damaged blood supply^[10, 11]^, abnormal lipid metabolism^[12, 13]^ and bone cell apoptosis^[14, 15]^ have been proposed to explain the occurrence and development of GONFH. However, the therapies developed according to the guideline established based on these theories have not been successful in prevention of GONFH progression. Clinical evidence about decreased replication and osteogenic differentiation capacity of bone marrow mesenchymal stromal cells (BMSCs)^[16–18]^ and the efficacy of BMSCs transplantation^[19–21]^ in GONFH patients indicate that GONFH might be a BMSCs-related disease. BMSCs contain mesenchymal progenitor cells which give rise to osteoblasts and adipocytes, and glucocorticoids have been shown to regulate osteogenic/adipogenic differentiation of BMSCs *in vitro*^[22, 23]^. Various treatments targeting on promoting osteogenic differentiation of BMSCs alleviate early necrotic phenotype in glucocorticoid-induced GONFH animal models^[24–26]^. Thus, we hypothesize that imbalanced osteogenic/adipogenic differentiation of BMSCs plays a dominant role in GONFH pathogenesis. Recent lineage tracing and single-cell RNA sequencing studies revealed that BMSCs are a group of heterogeneous multipotent cells^[27]^, and skeletal-derived mesenchymal progenitor cells including collagen II (Col2)^+^ lineage and Osterix (Osx)^+^ lineage compose a large proportion of BMSCs^[28, 29]^. However, which sub-population of BMSCs significantly contributes to the GONFH pathogenesis remains unknown.

BMSCs differentiation is a complex process and is regulated by multiple signaling pathways. Among them, canonical Wnt/β-catenin signaling functions as a switch in determining osteogenic/adipogenic differentiation of BMSCs^[30]^. When β-catenin is accumulated in the nucleus, interacting with TCF/Lef transcription factors to activate downstream target genes including Runt-related transcription factor 2 (Runx2) and Osterix^[31]^. On the contrary, inhibition of β-catenin induces expressions of CCAAT/enhancer binding protein alpha (C/EBPα) and peroxisome proliferator activated receptor gamma (PPAR-γ) for adipogenesis^[32, 33]^. Our previous study showed a significant decrease of β-catenin in the glucocorticoid-induced GONFH rat model^[34]^. Other groups also reported an involvement of β-catenin signaling in GONFH development^[35–39]^.

Here, abnormal osteogenesis and adipogenesis with decreased β-catenin signaling were observed in the necrotic femoral heads of GONFH patients and glucocorticoid-induced GONFH rats. Activation of β-catenin signaling effectively alleviated the necrotic changes in GONFH rats by restoring glucocorticoid exposure-induced imbalanced osteogenic/adipogenic differentiation of BMSCs. Interestingly, specific deletion of *β-catenin* in Col2^+^ cells, but not Osx^+^ cells led to a full spectrum of disease phenotype of GONFH in mice even including subchondral bone destruction and femoral head collapse. Our present study provided novel molecular mechanisms of GONFH pathogenesis, and an ideal genetic modified mouse model for GONFH study.

## Results

### Abnormal osteogenesis and adipogenesis with decreased β-catenin in necrotic femoral heads of GONFH patients

To determine the pathogenesis of GONFH, we harvested the surgical specimens from GONFH patients with femoral head necrosis (n=15) and samples from trauma patients without femoral head necrosis (n=10). Compared to the non-necrotic femoral heads, we observed subchondral bone destruction (Figure 1A, black asterisk), liquefied necrotic foci (Figure 1A, black arrow), deformed and collapsed outline in the necrotic femoral head. μCT analysis further showed sparse and cracked trabeculae in the collapsed region (Figure 1B, red box) and liquefied necrotic region (Figure 1B, yellow box) compared to the corresponding regions (Figure 1B, white boxes) in the non-necrotic femoral heads, which were further confirmed by the decreased BV/TV, Tb.N and Tb.Th and increased Tb.Sp (Figure 1C). The representative images of ABH staining (Figure 1D) revealed sparse trabeculae, massive accumulated fat droplets (Figure 1D, black triangles), numerous empty lacunae of osteocytes (Figure 1D, black arrows) and extensive subchondral bone destruction (Figure 1D, yellow arrows) in the necrotic femoral heads. Histomorphological quantitative analyses showed decreased trabecular bone area (Figure 1E), increased fat droplet area (Figure 1E) and empty lacunae rate (Figure 1F) in the necrotic femoral heads. TUNEL staining revealed more apoptotic osteocytes in the necrotic femoral heads (Figure 1G). We then detected the expression of osteogenic (Runx2, Osterix) and adipogenic (FABP4, PPAR-γ, CEBP/α) marker genes. The results of IHC staining showed a down-regulation of Runx2 and an up-regulation of FABP4 in the necrotic regions (Figure 1H). Western blot analysis showed increased expressions of PPAR-γ and CEBP/α and a decreased expression of Osterix in the human necrotic femoral head tissues (Figure 1I). These findings indicate an involvement of abnormal osteogenesis and adipogenesis during GONFH pathogenesis. β-catenin can regulate osteogenesis and adipogenesis downstream target genes (Runx2, Osterix, FABP4, PPAR-γ and CEBP/α) expressions^[40]^. The significant decrease of β-catenin in the necrotic femoral heads (Figure 1G, 1H) indicate an important role of β-catenin signaling in GONFH pathogenesis.

**Figure 1.**
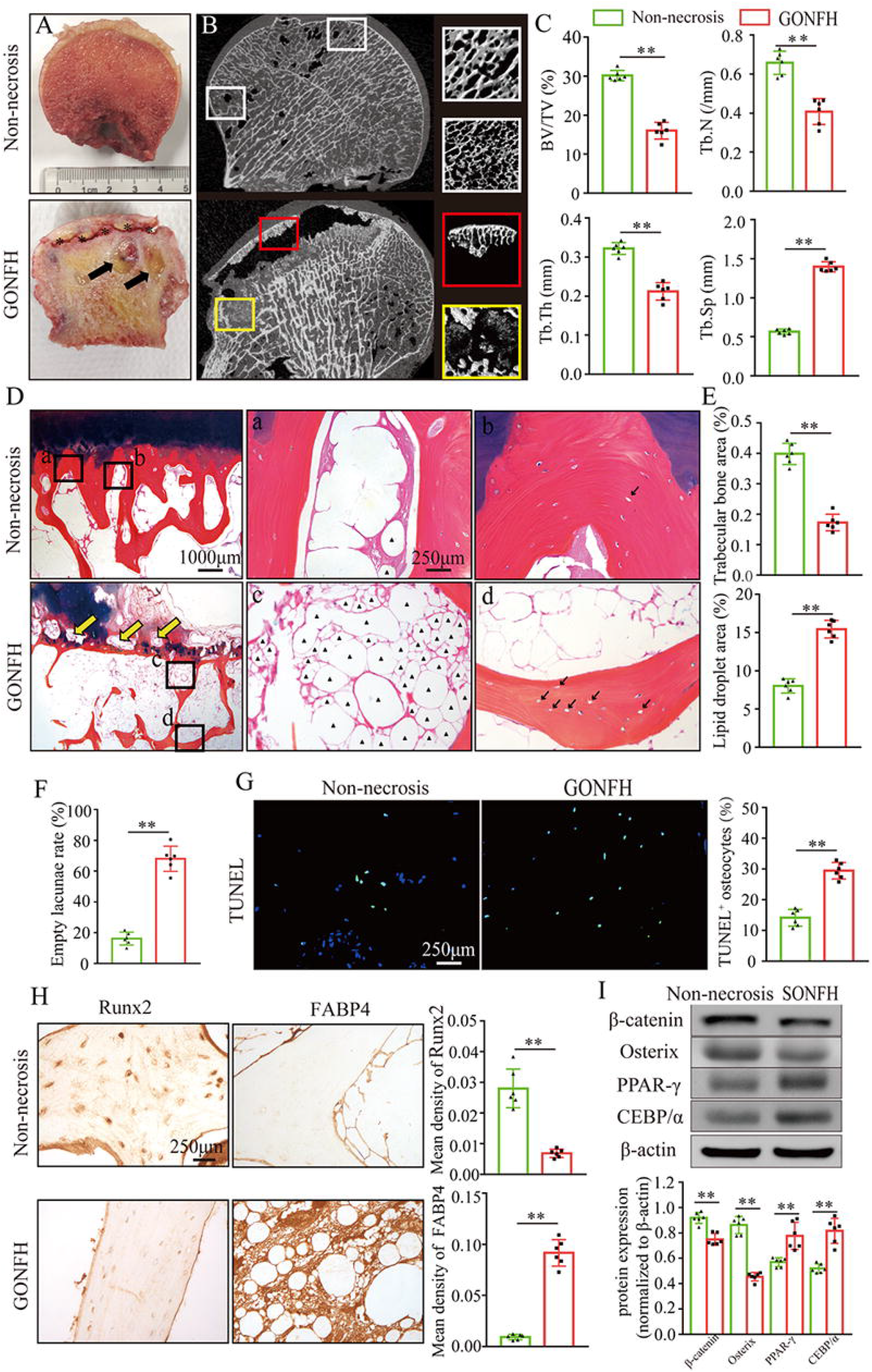
Abnormal osteogenesis and adipogenesis with decreased β-catenin signaling in the necrotic femoral heads of GONFH patients. Necrotic (n=15) and Non-necrotic (n=10) femoral head samples were obtained from the patients with GONFH or femoral neck fracture, respectively. (A) Gross anatomy analysis of human necrotic and non-necrotic femoral head samples. Black asterisks: subchondral collapsed region. Black arrows: liquefied necrotic region. (B) μCT images of human necrotic and non-necrotic femoral heads. Red and Yellow boxed area: 3D images of subchondral collapsed region and liquefied necrotic region, respectively. White boxed areas: 3D images of corresponding regions in the non-necrotic femoral heads. (C) Quantitative analysis of BV/TV, Tb.N, Tb.Th and Tb.Sp on the necrotic regions. (D) ABH staining of human necrotic and non-necrotic femoral heads. a,c: high magnification images of bone marrow; b,d: high magnification images of bone trabeculae; Yellow arrows: subchondral bone destruction; black arrows: empty lacunae of osteocytes; black triangles: fat droplets. (E) Histomorphological quantitative analysis of trabecular bone area and fat droplet area. (F) Histomorphological quantitative analysis of empty lacunae rate. (G) TUNEL staining of osteocytes in human necrotic and non-necrotic femoral heads. (H) IHC staining of Runx2 and FABP4 expressions in human necrotic and non-necrotic femoral heads. (I) Western blot of β-catenin, Osterix, PPAR-γ and CEBP/α in human necrotic and non-necrotic femoral heads.

### Inhibition of β-catenin signialing leads to abnormal osteogenesis and adipogenesis in glucocorticoid-induced GONFH rats

To further analyze the role of β-catenin in GONFH pathogenesis, a glucocorticoid-induced GONFH rat model was established by continuous MPS induction, as previously described^[9]^. Gross anatomy analysis showed a local melanocratic region on the femoral head surface of GONFH rats without any structure deformity or collapse (Figure 2A, black arrow). No color change was observed on the femoral head surface from the Wnt agonist 1 treated rats. μCT images and quantitative analysis of bone microstructure showed severe trabecular bone loss in the femoral heads of GONFH rats, especially in the subchondral region (Figure 2A, 2B). Systemic injection of Wnt agonist 1 effectively increased bone mass (Figure 2A), and alleviated decreased BV/TV, Tb.N and Tb.Th and increased Tb.Sp in GONFH rats (Figure 2B). ABH staining and histomorphological quantitative analysis revealed that the GONFH rats presented sparse trabeculae, massive fat accumulation (Figure 2C, black arrow) and numerous empty lacunae of osteocytes (Figure 2C, black arrows) in the necrotic region of femoral heads (Figure 2C-2E). However, no subchondral bone destruction, femoral head collapse and deformity was occurred in this GONFH rat model (Figure 2C). TUNEL staining revealed increased apoptotic osteocytes in the femoral heads of GONFH rats (Figure 2F). ALP is an osteoblast marker reflecting the capacity of osteogenesis. FABP4 is essential for lipid formation and metabolism. The decreased β-catenin and ALP expressions and increased FABP4 expression confirmed abnormal osteogenesis and adipogenesis during rat GONFH development (Figure 2G, 2H), consistent with the findings in human GONFH femoral heads. Interestingly, activation of β-catenin by systematically injecting Wnt agonist 1 for 6 weeks attenuated the early necrotic changes (Figure 2C-2E) and osteocyte apoptosis (Figure 2F), and restored ALP and FABP4 expressions in the femoral heads of GONFH rats (Figure 2G, 2H). Overall, these data indicate that β-catenin inhibition-induced abnormal osteogenesis and adipogenesis contributes to GONFH pathogenesis.

**Figure 2.**
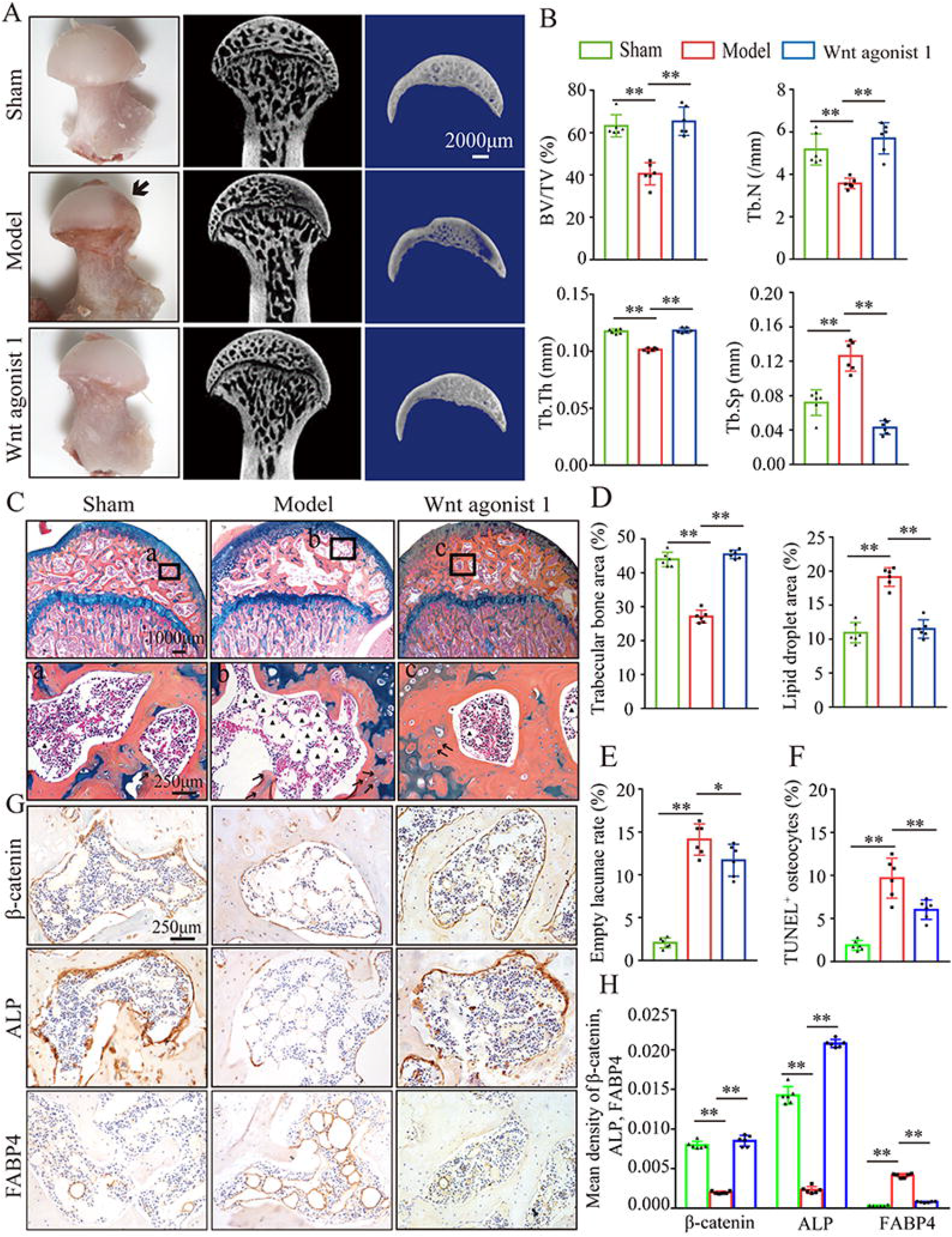
Systemic injection of Wnt agonist 1 alleviates abnormal osteogenesis and adipogenesis in rat GONFH femoral heads. The rat model of GONFH was established by a co-induction of lipopolysaccharide and methylprednisolone (MPS). The femoral head samples were harvested after GONFH rats were intravenously injected with Wnt agonist 1 for 6 weeks. (A) Gross anatomy analysis and μCT images of femoral heads in sham, model and Wnt agonist 1 treated groups. (B) μCT quantitative analysis of BV/TV, Tb.N, Tb.Th and Tb.Sp in each group. (C) ABH staining of femoral heads in each group. a-c, high magnification images of the representative region; black arrows: empty lacunae of osteocytes; black triangles: fat droplets. (D) Histomorphological quantitative analysis of trabecular bone area and fat droplet area in each group. (E) Histomorphological quantitative analysis of empty lacunae rate in each group. (F) TUNEL staining of osteocytes in the femoral heads of GONFH rats. (G, H) IHC staining and quantitative analysis of β-catenin, ALP and FABP4 in rat femoral heads of each group.

### β-catenin directs dexamethasone-induced osteogenic/adipogenic differentiation of rat BMSCs

We further investigated cellular mechanisms involved in the necrotic femoral heads of glucocorticoid-induced GONFH rats. Primary bone marrow cells were extracted from the proximal femur of 4-week-old SD rats and cultivated to P3. Flow cytometry analysis identified that they were BMSCs, with CD90^+^, CD29^+^, CD45^−^ and CD11b^−^ characteristics (Figure 3A). ALP staining showed that the osteogenic differentiation of BMSCs were significantly inhibited after exposing to 10^2^ nM or higher concentration of Dex for 7 days (Figure 3B). Oil red O staining showed a progressive adipogenic differentiation of BMSCs with the increased concentration of Dex (Figure 3B). These findings indicated that increased adipogenic differentiation and decreased osteogenic differentiation of BMSCs caused the abnormal osteogenesis and adipogenesis, finally leading to the femoral head necrosis in GONFH rats. The decreased expressions of β-catenin, Runx2, ALP and increased expressions of PPAR-γ, CEBP/α further revealed that Dex exposure inhibited β-catenin signaling, and redirected BMSCs differentiation from osteoblasts to adipocytes (Figure 3C, 3D). However, this Dex-induced abnormal differentiation of BMSCs were restored by SKL2001 through activation of β-catenin signaling (Figure 3E-3G). Taken together, these findings suggest that β-catenin inhibition redirects the direction of BMSCs differentiation, contributing to the GONFH development.

**Figure 3.**
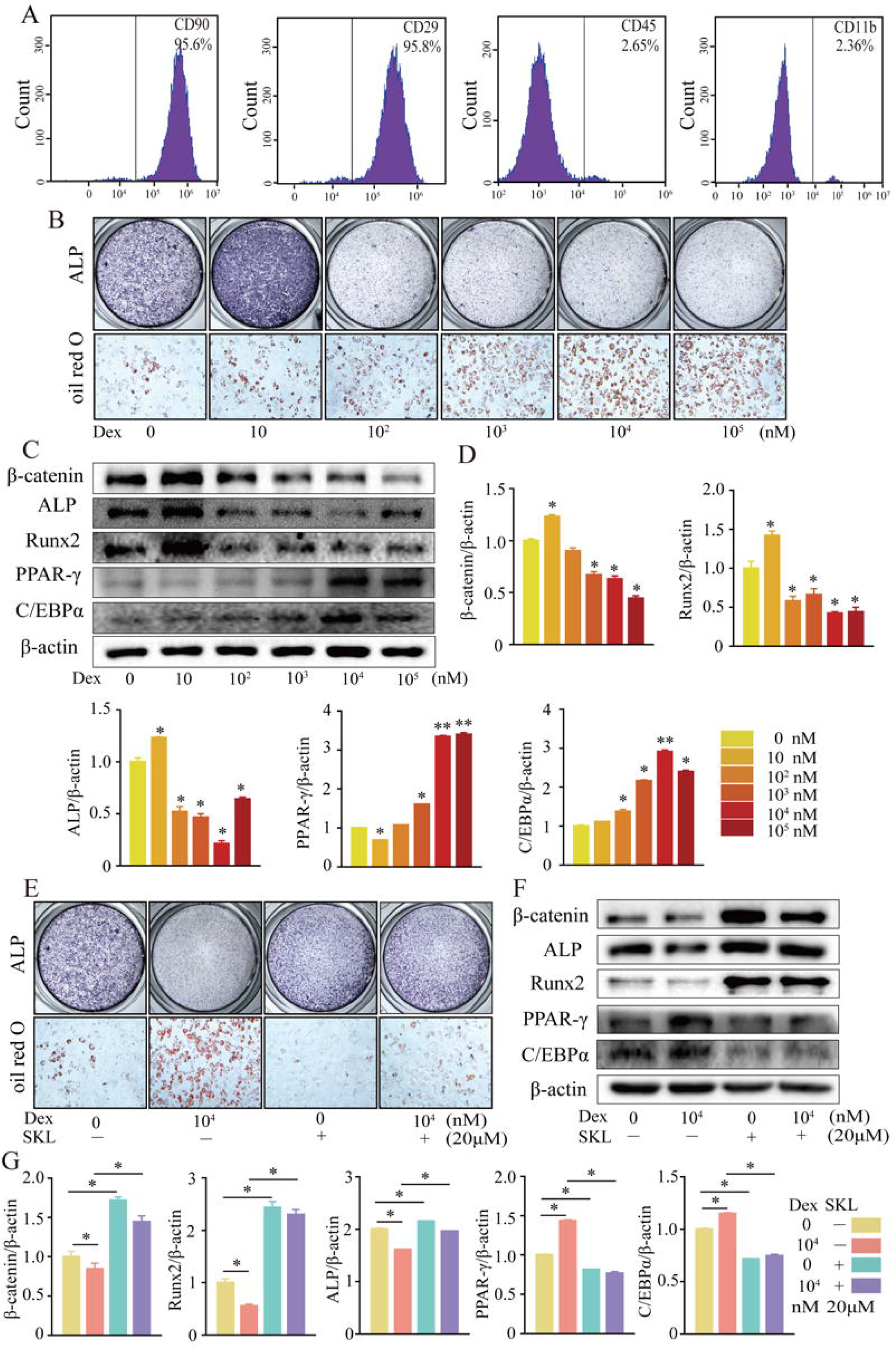
Activation of β-catenin redirected Dex-induced imbalanced osteogenic/adipogenic differentiation of BMSCs. Primary BMSCs were extracted from the proximal femur of 4-week-old SD rats and cultivated to passage 3 for subsequent experiments. (A) Flow cytometry analyzing the surface markers of rat BMSCs. (B) ALP staining and oil red O staining of BMSCs at the increasing concentrations of dexamethasone (Dex). (C, D) Western blot and quantitative analysis of β-catenin, Runx2, ALP, PPAR-γ and CEBP/α in BMSCs at the increasing concentrations of Dex. (E) ALP staining and oil red O staining of rat BMSCs at the condition with or without 10^4^ nM Dex and 20 μM SKL2001 (an agonist of β-catenin). (F, G) Western blot and quantitative analysis of β-catenin, Runx2, ALP, PPAR-γ and CEBP/α in rat BMSCs at the condition with or without 10^4^ nM Dex and 20 μM SKL2001.

### Loss-of-function of β-catenin in Col2^+^, but not Osx^+^, -expressing cells leads to a GONFH-like phenotype in femoral head

Skeletal progenitor cells are major source of BMSCs, and both Col2^+^ lineage and Osx^+^ lineage can trans-differentiate into BMSCs^[27, 28]^. To determine which subset of BMSCs contributes more to GONFH pathogenesis, we generated *β-catenin^Col2ER^* mice and *β-catenin^OsxER^* mice, in which *β-catenin* gene was specifically deleted in Col2^+^ cells and Osx^+^ cells, respectively. For mapping their cell fate in the femoral head, we induced *Tomato^Col2ER^* mice and *Tomato^OsxER^* mice at 2-weeks old and analyzed at the age of 1 month. The fluorescent images showed that Col2^+^ cells were evidently expressed in the chondrocytes of articular cartilage, secondary ossification center and growth plate (Figure 4A, white arrowheads), osteoblasts (Figure 4A, green arrowheads), osteocytes (Figure 4A, yellow arrowheads) and stromal cells (Figure 4A, red arrowheads) underneath the growth plate, while Osx^+^ cells were identified only in osteoblasts, osteocytes and stromal cells underneath the growth plate (Figure 4A). We then trace Col2^+^ lineage for 9 months and found that Col2^+^ chondrocytes in secondary ossification center and growth plate were replaced by Col2^+^ osteoblasts, osteocytes and stromal cells with age (Figure 4B), and these Col2^+^ cells continuously produced osteoblasts and osteocytes in the femoral head (Figure 4B). These findings indicated the self-renew ability and osteogenic commitment of Col2^+^ cells. We also induced *Tomato^Col2ER^* mice at the age of 1 month, but identified a poor *Cre*-recombinase efficiency 24 hours and 3 months later (Figure 4C). IHC assay showed a significant decrease of β-catenin in the femoral heads of 3-month-old *β-catenin^Col2ER^* mice and *β-catenin^OsxER^* mice (Figure 4D, 4E), indicating successful establishments of these *β-catenin* conditional knockout mouse models.

**Figure 4.**
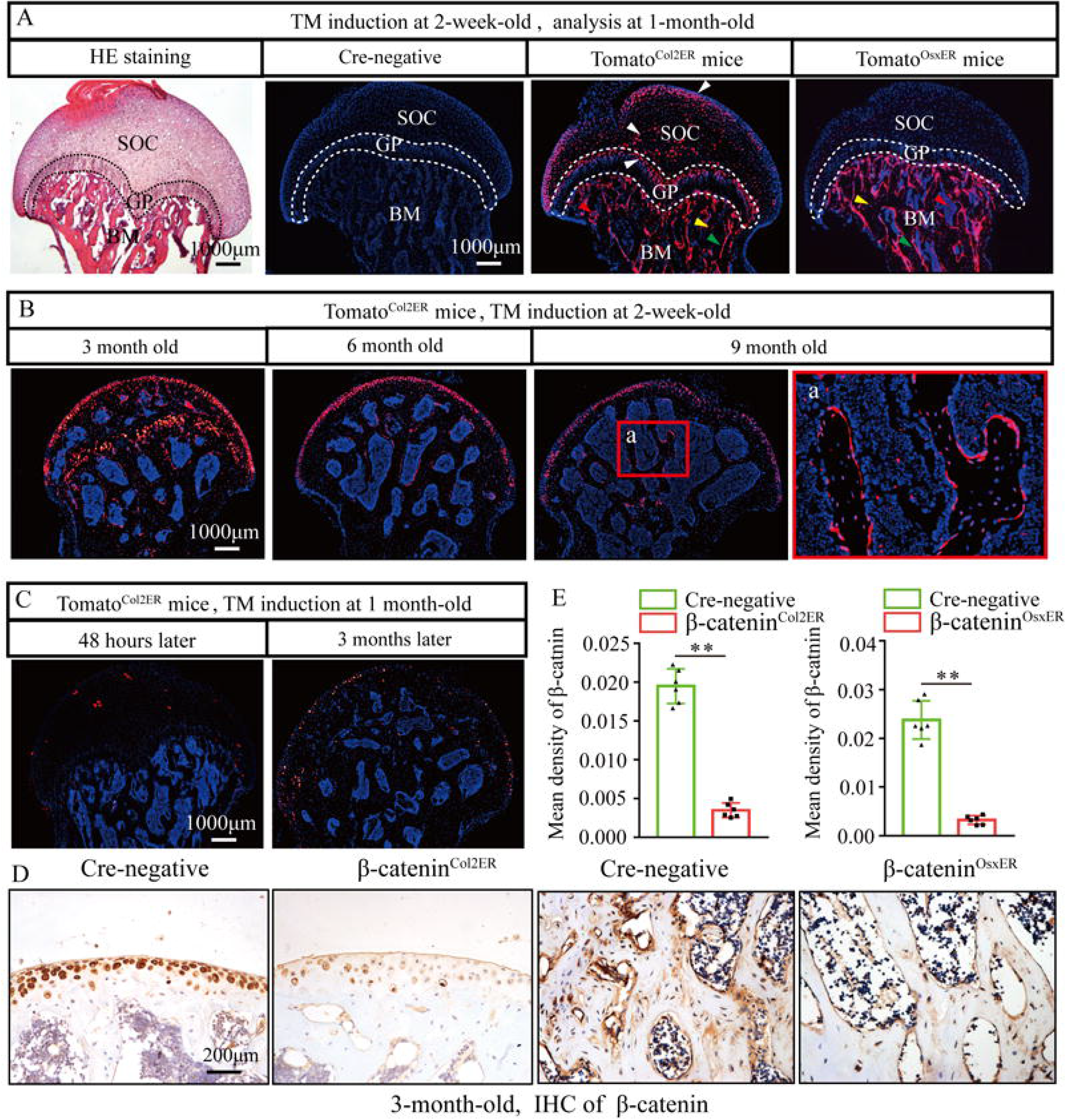
Fate mapping of Col2^+^ cells and Osx^+^ cells in the femoral head. *Tomato^Col2ER^* mice and *Tomato^OsxER^* mice were continuously received 5 dose of TM injections (1 mg/10 g body weight) at the age of 2 weeks for fate mapping analysis. (A) Distributions of Col2^+^ and Osx^+^ cells in the femoral heads of 1-month-old *Tomato^Col2ER^* mice and *Tomato^OsxER^*mice. White arrowheads: chondrocytes; Green arrowheads: osteoblasts; Yellow arrowheads: osteocytes; Red arrowheads: bone marrow stromal cells; GP: growth plate; SOC: second ossification center; BM: bone marrow. (B) Lineage trace of Col2^+^ cells in the femoral head for 9 months. a: high magnification image of bone marrow region. (C) Poor *Cre*-recombinase efficiency in the femoral heads of *Tomato^Col2ER^* mice with TM induction at the age of 1 month. (D, E) Femoral heads were harvested from 3-month-old *β-catenin^Col2ER^* mice and *β-catenin^OsxER^* mice to detect expression of βLcatenin. IHC staining and quantitative analysis of β-catenin in the femoral heads of 3-month-old *β-catenin^Col2ER^* mice and *β-catenin^OsxER^* mice.

Then, we analyzed morphologic changes of the femoral heads from 3-months old *β-catenin^Col2ER^*mice and *β-catenin^OsxER^*mice. No change of appearance was observed on the femoral heads both in *β-catenin^Col2ER^* mice and *β-catenin^OsxER^*mice compared to *Cre*-negative littermates (Figure 5A, 6A). μCT images and quantitative analysis of decreased BV/TV, Tb.N, Tb.Th and increased Tb.Sp indicated that both *β-catenin^Col2ER^*mice and *β-catenin^OsxER^*mice presented severe bone loss in the femoral heads (Figure 5B, 6B). ABH staining and histomorphological analysis further revealed that bone trabeculae were extensively replaced by fat droplet clusters in the femoral heads (especially in the subchondral region) of *β-catenin^Col2ER^* mice (Figure 5C, 5D), similar to the necrotic femoral heads in GONFH patients and glucocorticoid-induced rat models. However, no fat droplet accumulation but only sparse and thin bone trabeculae were observed in the femoral heads of *β-catenin^OsxER^*mice (Figure 6C, 6D). Compared to *Cre*-negative littermates, the number of empty lacunae was increased in the femoral heads in *β-catenin^Col2ER^*mice (Figure 5E), but no changed in *β-catenin^OsxER^*mice (Figure 6E). TUNEL staining revealed more apoptotic osteocytes in *β-catenin^Col2ER^* mice compared to *Cre*-negative littermates (Figure 5F). All these findings indicate that loss-of-function of β-catenin in Col2^+^ cells, but not in Osx^+^ cells, led to an GONFH-like phenotype in femoral head. Furthermore, decreased Runx2, ALP and increased PPAR-γ, CEBP/α expressions in *β-catenin^Col2ER^* mice (Figure 5G) revealed that this GONFH-like phenotype was caused by imbalance of osteogenic/adipogenic differentiation of Col2^+^ cells.

**Figure 5.**
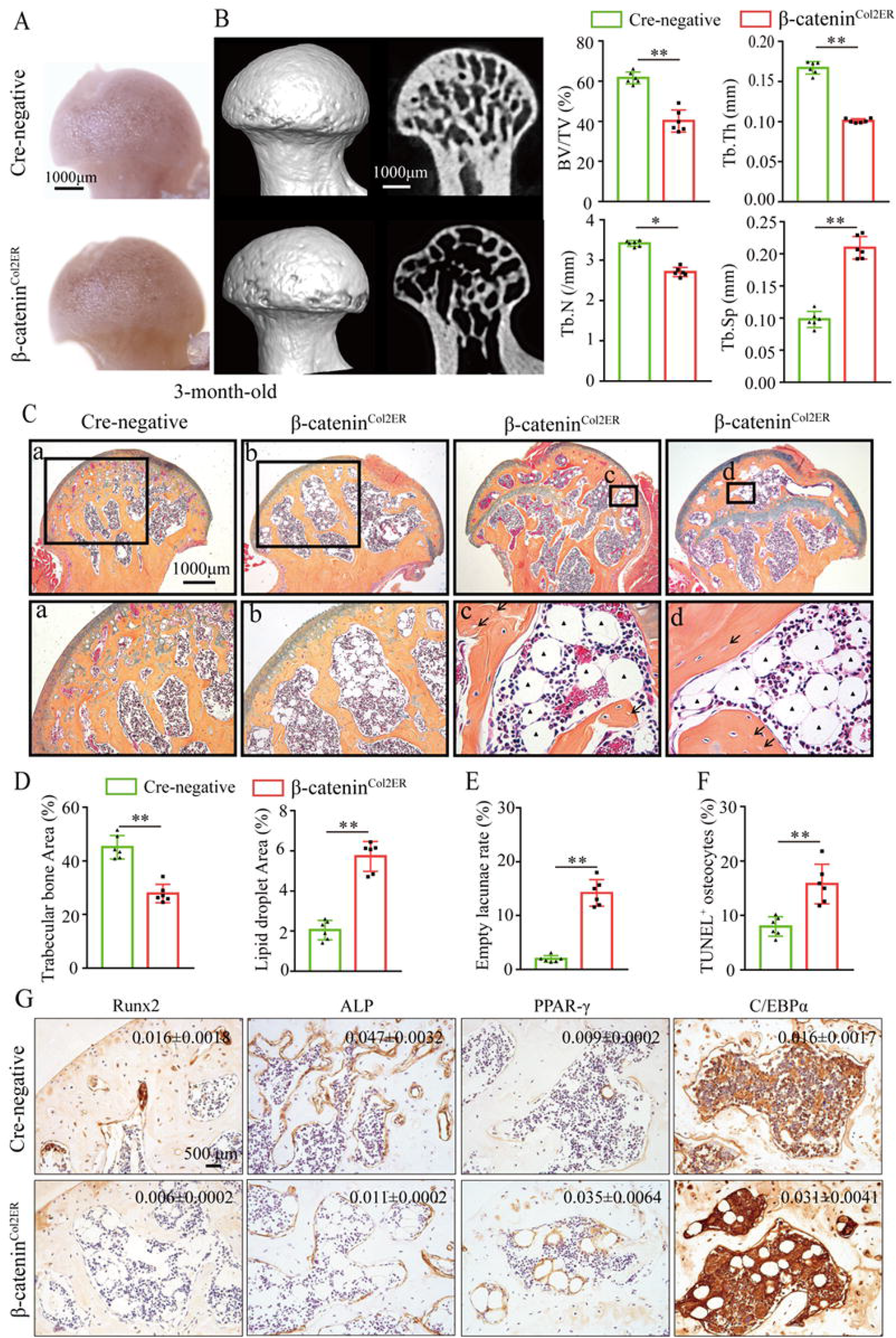
Deletion of *β-catenin* in Col2^+^ cells leads to a GONFH-like phenotype. Femoral heads were harvested from 3-month-old *β-catenin^Col2ER^*mice and *Cre-*negative littermates with 5 continuous dosage of TM injections (1 mg/10 g body weight) at the age of 2 weeks. (A) Gross anatomy analysis of femoral heads in *β-catenin^Col2ER^*mice and *Cre-*negative littermates. (B) Representative μCT images and quantitative analysis of BV/TV, Tb. N, Tb. Th and Tb. Sp in the femoral heads of *β-catenin^Col2ER^* mice. (C) ABH staining of femoral heads in *β-catenin^Col2ER^* mice. a-d: high magnification images of representative subchondral bone region; Black triangle arrowheads: fat droplets; Black arrows: empty lacunae of osteocytes. (D) Histomorphological quantitative analysis of trabecular bone area and fat droplet area in the femoral heads of *β-catenin^Col2ER^* mice. (E) Histomorphological quantitative analysis of empty lacunae rate in the femoral heads of *β-catenin^Col2ER^*mice. (F) TUNEL staining of osteocytes in the femoral heads of *β-catenin^Col2ER^* mice. (G) IHC staining and quantitative analysis of Runx2, ALP, PPAR-γ and CEBP/α in the femoral heads of *β-catenin^Col2ER^* mice.

**Figure 6.**
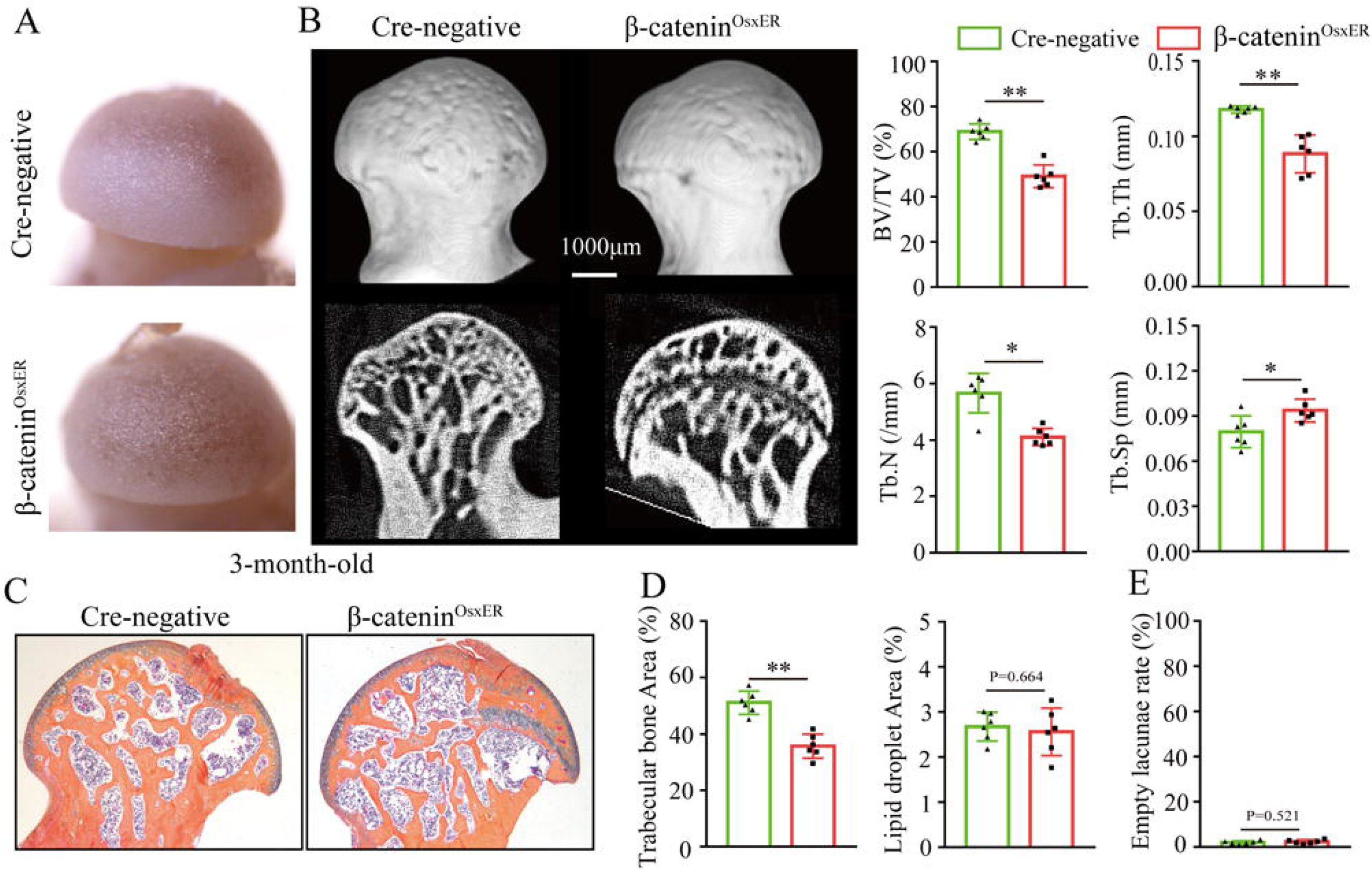
Loss-of-function of *β-catenin* in Osx^+^ cells causes bone loss in the femoral heads. Femoral heads were harvested from 3-month-old *β-catenin^OsxER^* mice and *Cre-*negative littermates with 5 continuous dosage of TM injections (1 mg/10 g body weight) at the age of 2 weeks. (A) Gross anatomy analysis of femoral heads in *β-catenin^OsxER^* mice and *Cre-*negative littermates. (B) μCT images and quantitative analysis of BV/TV, Tb. N, Tb. Th and Tb. Sp in the femoral heads of *β-catenin^OsxER^* mice. (C) ABH staining of femoral heads in *β-catenin^OsxER^*mice. (D) Histomorphological quantitative analysis of trabecular bone area and fat droplet area in the femoral heads of *β-catenin^OsxER^*mice. (E) Histomorphological quantitative analysis of empty lacunae rate in the femoral heads of *β-catenin^OsxER^*mice.

### Older *β-catenin^Col2ER^*mice display subchondral bone destruction and collapse tendency in the femoral heads

Femoral head collapse occurs at the end-stage of GONFH and represents a poor joint function^[41]^. To determine the occurrence of femoral head collapse, we further analyzed 6-month-old *β-catenin^Col2ER^* mice. Gross appearance showed that older *β-catenin^Col2ER^* mice lost smooth and regular outline on the femoral heads compared to *Cre*-negative littermates and 3-month-old *β-catenin^Col2ER^*mice (Figure 7A). μCT images showed severe bone loss (Figure 7B, 7C), subchondral bone destruction (Figure 7B, yellow arrows), local collapse (Figure 7B, red arrows) and integral deformity (Figure 7B, red dotted lines) in the femoral heads of older *β-catenin^Col2ER^* mice. μCT quantitative analysis confirmed a significant increase of subchondral bone defect area on the femoral head surface in older *β-catenin^Col2ER^*mice (Figure 7D). Besides of these early necrotic changes including sparse bone trabeculae, accumulated fat droplets (Figure 7E, black triangles) and increased empty lacunae of osteoctyes (Figure 7E, black arrows, Figure 7F), ABH staining also showed subchondral bone destruction (Figure 7E, yellow arrows), femoral head collapse or deformity (Figure 7E, red arrows and red dotted lines) in older *β-catenin^Col2ER^*mice. Compared to *Cre*-negative littermates, the older *β-catenin^Col2ER^* mice presented a decrease of loading-bearing stiffness, indicating a poor biomechanical support of the femoral heads (Figure 7G).

**Figure 7.**
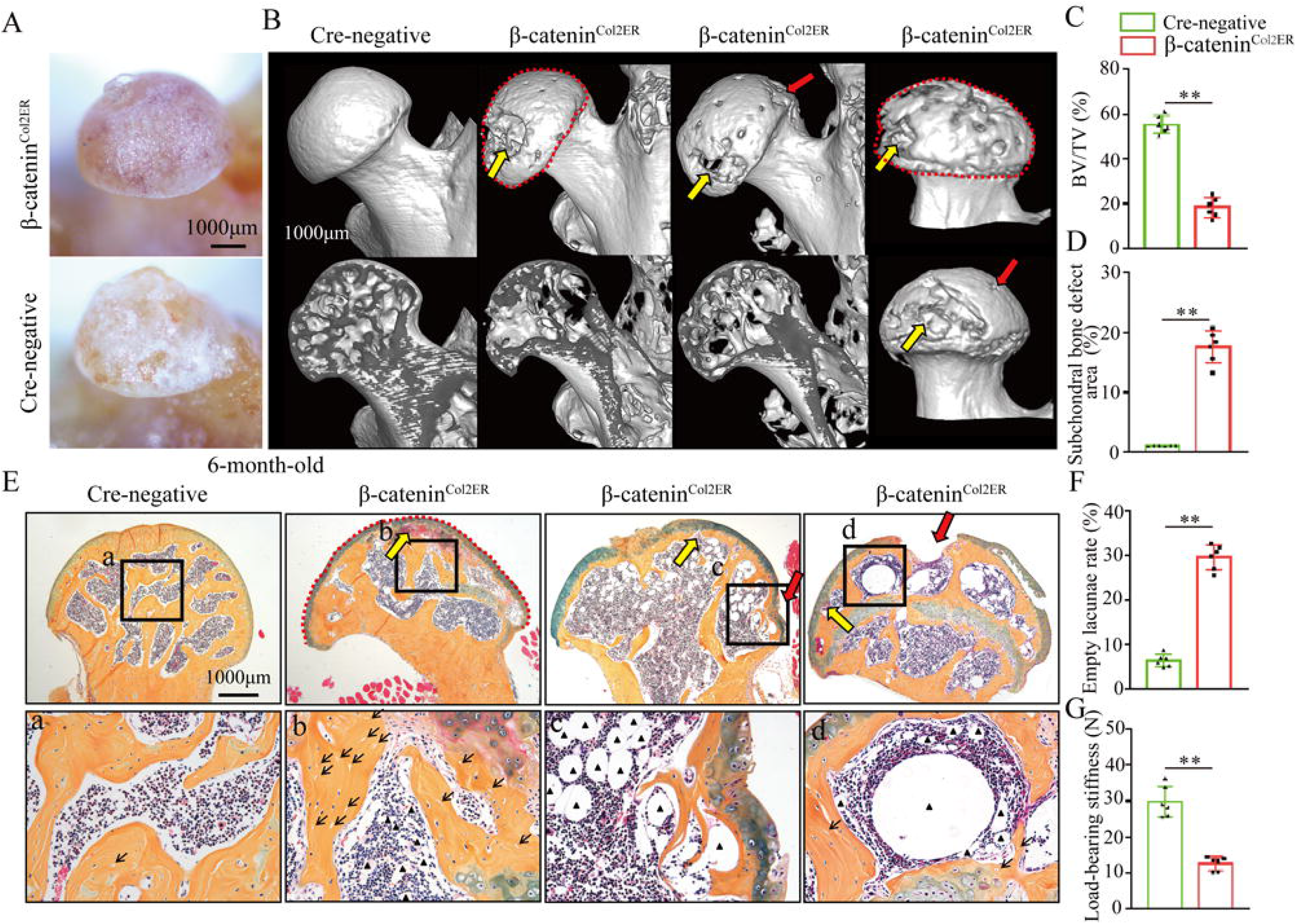
Older *β-catenin^Col2ER^* mice display subchondral bone destruction and collapse tendency in femoral heads. Femoral heads were harvested from 6-month-old *β-catenin^Col2ER^* mice and *Cre-*negative littermates with 5 continuous dosage of TM injections (1 mg/10 g body weight) at the age of 2 weeks. (A) Gross anatomy analysis of femoral heads in 6-month-old *β-catenin^Col2ER^*mice. (B) Representative μCT images of femoral heads in 6-month-old *β-catenin^Col2ER^*mice. Red dotted lines: integral deformity. Red arrows: local collapse. Yellow arrows: subchondral bone destruction. (C) Quantitative analysis of BV/TV in the femoral heads in 6-month-old *β-catenin^Col2ER^*mice. (D) Quantitative analysis of subchondral bone defect area on the femoral head surface in older *β-catenin^Col2ER^* mice. (E) ABH staining of femoral heads in 6-month-old *β-catenin^Col2ER^* mice. a-c: high magnification images of representative subchondral bone region. Red dotted lines: integral deformity. Red arrows: local collapse. Black arrows: empty lacunae of osteoctyes. Black triangles: fat droplets. (F) Histomorphological quantitative analysis of empty lacunae rate. (G) Loading-bearing stiffness of the femoral heads in 6-month-old *β-catenin^Col2ER^*mice.

### Wnt agonist 1 could not alleviate GONFH-like phenotype in *β-catenin^Col2ER^*mice

As the therapeutic effects of Wnt agonist 1 on rat GONFH, we further determined whether it could alleviate the GONFH-like phenotype in *β-catenin^Col2ER^* mice. After induction with TM, *β-catenin^Col2ER^*mice were immediately injected with Wnt agonist 1 three times once week until they were analyzed at the age of 3 months. However, μCT images and quantitative analysis of BV/TV, Tb.N, Tb.Th and Tb.Sp showed severe bone loss were not restored in *β-catenin^Col2ER^* mice by treating with Wnt agonist 1 (Figure 8A, 8B). Accumulated fat droplets, sparse trabeculae and empty lacunae were still existed in the femoral heads of Wnt agonist 1 treated *β-catenin^Col2ER^* mice (Figure 8C-8E). IHC staining showed no change of decreased ALP and increased FABP4 in the femoral heads of *β-catenin^Col2ER^*mice after Wnt agonist 1 treatment (Figure 8F, 8G). These findings indicate that Wnt agonist 1 could not restore the imbalanced osteogenic/adipogenic differentiation of Col2^+^ cells in *β-catenin^Col2ER^*mice.

**Figure 8.**
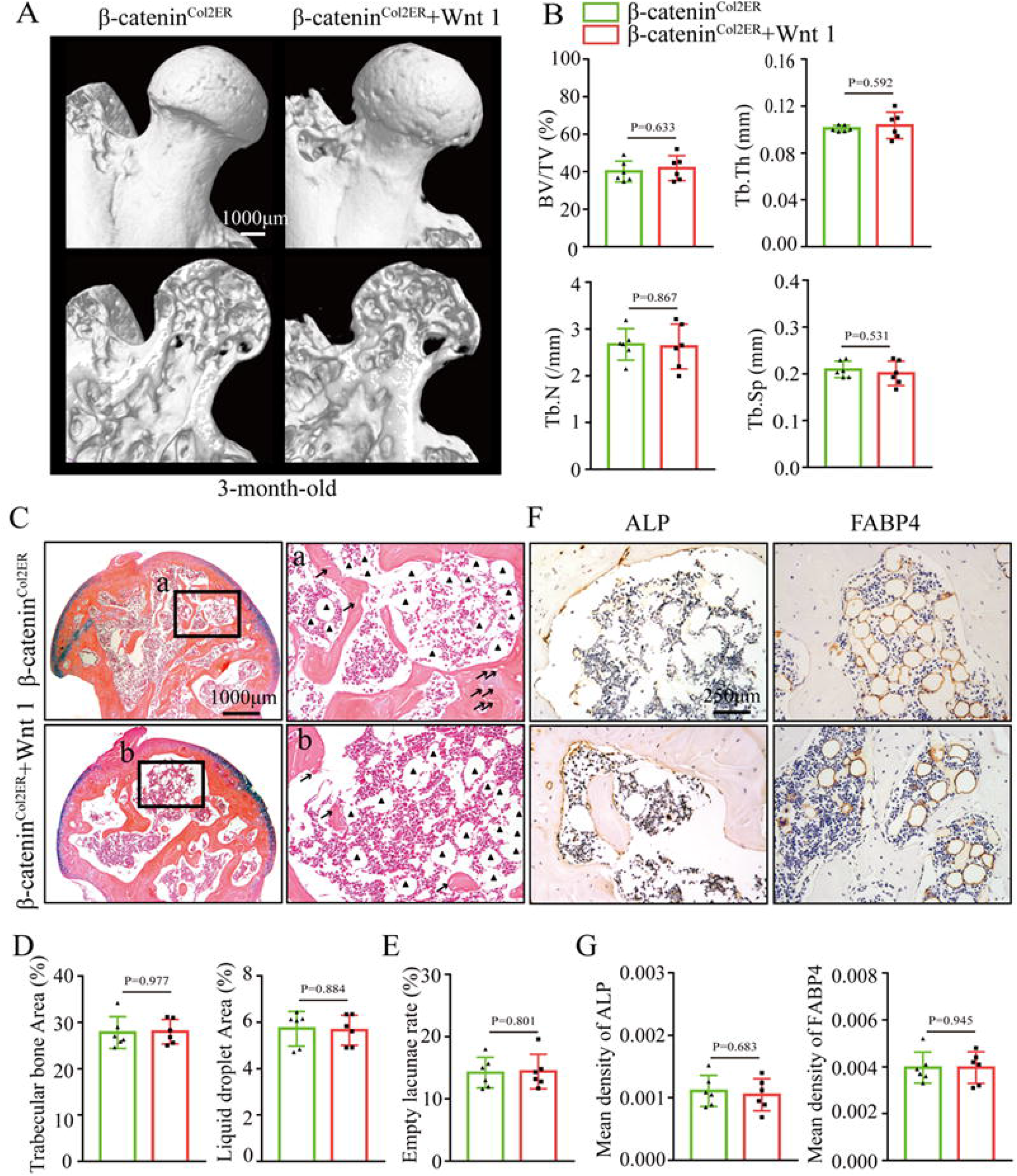
Systemic injection of Wnt agonist 1 can not alleviate the GONFH-like phenotype in *β-catenin^Col2ER^* mice. *β-catenin^Col2ER^* mice with 5 continuous dosage of TM injections (1 mg/10 g body weight) were treated with Wnt agonist 1 three times once week until sacrifice at the age of 3 months. (A) Representative μCT images of femoral heads in Wnt agonist 1 treated *β-catenin^Col2ER^* mice. (B) Quantitative analysis of BV/TV, Tb.N, Tb.Th and Tb.Sp in the femoral heads of Wnt agonist 1 treated *β-catenin^Col2ER^*mice. (C) ABH staining of the femoral heads in Wnt agonist 1 treated *β-catenin^Col2ER^* mice. a-b: high magnification images of subchondral bone region. Black triangles: fat droplets. (D) Histomorphological quantitative analysis of trabecular bone area and fat droplet area in the femoral heads of Wnt agonist 1 treated *β-catenin^Col2ER^* mice. (E) Histomorphological quantitative analysis of empty lacunae rate in the femoral heads of Wnt agonist 1 treated *β-catenin^Col2ER^*mice. (F, G) IHC staining and quantitative analysis of ALP and FABP4 in the femoral heads of Wnt agonist 1 treated *β-catenin^Col2ER^*mice.

## Discussion

Despite it is well known that glucocorticoids are the certain etiology of GONFH, its pathogenesis remains largely unknown. Current evidence including reduced osteogenic differentiation ability of BMSCs from GONFH patients and clinical therapeutical effects of BMSCs transplantation indicates that GONFH could be resulted from abnormal differentiation of BMSCs^[18–21]^. Glucocorticoid-induced animal models are commonly used for GONFH research, and several previous studies have revealed that direct BMSCs modifications with miRNA^[42, 43]^, lncRNA^[44, 45]^ and circRNA^[24, 46]^ or microenvironment interventions by injecting platelet-rich plasma^[47, 48]^, exosome^[49, 50]^ and growth factors^[51, 52]^ could promote osteogenic differentiation of BMSCs and exhibit anti-GONFH effects in animals. In the present studies, we found severe trabecular bone loss and massive fat droplets accumulation in the necrotic femoral heads of GONFH patients and glucocorticoid-induced GONFH rats, confirming increased adipogenesis and decreased osteogenesis in GONFH pathogenesis. Dex inhibited osteogenic differentiation of rat BMSCs and promoted their adipogenic differentiation, revealing that this abnormal osteogenesis and adipogenesis in GONFH rats were caused by the imbalance in osteogenic/adipogenic differentiation of BMSCs. As β-catenin plays an important role in BMSCs differentiation, we then analyzed its role in GONFH pathogenesis. The results showed a significant inhibition of β-catenin signaling in the necrotic femoral heads of GONFH patients and glucocorticoid-induced GONFH rats, while Wnt agonist 1 attenuated accumulated fat droplets and sparse trabeculae in GONFH rats by activation of β-catenin signaling. The subsequent cellular experiments revealed that β-catenin activation restored the imbalance of osteogenic/adipogenic differentiation of BMSCs caused by long-term exposure of Dex. Overall, these findings indicate that β-catenin inhibition-induced imbalanced osteogenic/adipogenic differentiation of BMSCs contributed to GONFH.

BMSCs are a group of heterogeneous cells, and their majorities are derived from skeletal progenitors^[27]^. It has reported that both Col2^+^ progenitors and Osx^+^ progenitors provide a large proportion of BMSCs^[28]^. Adult transgenic mice with conditional deletion of *β-catenin* in Col2^+^ cells, but not Osx^+^ cells, displayed a GONFH-like phenotype. Extensive trabecular bone were replaced by fat droplets in the femoral head (especially in the subchondral bone region) of *β-catenin^Col2ER^* mice compared to *β-catenin^OsxER^* mice. Cell fate mapping revealed a apparent difference in the distributions of Osx^+^ cells and Col2^+^ cells in the femoral head when mice induced at 2-week-old. Besides of the similar parts with Osx^+^ cells (osteoblasts, osteocytes and stromal cells underneath growth plate), Col2^+^ cells also largely expressed in the chondrocytes of articular cartilage, secondary ossification center and growth plate. Whether these Col2^+^ chondrocytes in secondary ossification center and growth plate caused this GONFH-like phenotype, as it has been reported that the osteogenic commitment of these Col2^+^ chondrocytes^[53, 54]^. At the age of 3 month, Col2^+^ chondrocytes in the secondary ossification center and growth plate were almost replaced by Col2^+^ osteoblasts, osteocytes and stromal cells, while GONFH-like phenotype including fat droplets accumulation and sparse bone trabeculae could be still observed in 6-month-old β-catenin^Col2ER^ mice, indicating that not only Col2^+^ chondrocytes but all Col2^+^ cells in different cellular morphology were involved in this GONFH-like phenotype. Nine months of lineage trace further revealed that Col2^+^ cells had self-renew ability and could continuously produced osteoblasts. The decreased ALP, Runx2 and increased CEBP/α and PPAR-γ in *β-catenin^Col2ER^* mice revealed that deletion of *β-catenin* shifted the commitment of Col2^+^ cells from osteoblasts to adipocytes, which should be responsible for this GONFH-like phenotype.

Femoral head collapse is the key feature of late stage GONFH and is disastrous to hip joint^[55]^. Although it has been well recognized that poor mechanical supports caused structural collapse of necrotic femoral head^[56]^, its detailed pathogenesis is poorly understood due to lack of ideal animal models. Glucocorticoid-induced bipedal and quadrupedal animal models only exhibit the early necrotic changes of GONFH^[57]^, without structure collapse of femoral head in MPS-induced rats. Other models using chemical or physical approaches are all failed to mimic full-range osteonecrosis of GONFH in humans^[58]^. Importantly, we found that femoral head collapse occurred spontaneously in older *β-catenin^Col2ER^* mice. Besides the early necrotic changes including trabecular bone loss, empty lacunae and fat droplets accumulation, extensive subchondral bone destruction were identified in 6-month-old *β-catenin^Col2ER^* mice. Previous studies have revealed that strong trabeculae and integrated subchondral bone are critical to maintain femoral head morphology^[59, 60]^. A significant decrease of loading-bearing stiffness confirmed a poor biomechanical supports in 6-month-old *β-catenin^Col2ER^* mice. Therefore, it could be concluded that β-catenin inhibition-induced imbalanced osteogenic/adipogenic differentiation of Col2^+^ cells caused a poor biomechanical supports (manifesting with sparse trabeculae and subchondral bone destruction), eventually leading to femoral head collapse in *β-catenin^Col2ER^* mice.

Osteocytes are the bone cells with longest life, up to decades within their mineralized environment^[61, 62]^. However, numerous empty lacunae of osteocyte were observed in the necrotic femoral head of GONFH patients and glucocorticoid-induced rats. Glucocorticoids are known to induce cell apoptosis^[63]^, and TUNEL staining confirmed increased osteocyte apoptosis in the human and rat necrotic femoral heads, leaving behinds these empty lacunae. The importance of β-catenin in the regulation of apoptosis is well established in skeletal system^[64, 65]^. Consistently, the expression of β-catenin was significantly decreased in the human and rat necrotic femoral heads. Moreover, we found that genetic deletion of β-catenin in Col2^+^ cells spontaneously caused osteocyte apoptosis in mice and activation of β-catenin by systematically injecting Wnt agonist 1 alleviated osteocyte apoptosis in glucocorticoid-induced rats, which indicate that β-catenin inhibition-induced osteocyte apoptosis contributed to empty lacunae of GONFH.

There are some limitations in the present study. Firstly, Wnt agonist 1 is a cell-permeating activator of Wnt signaling that induces transcriptional activity dependent on β-catenin^[66, 67]^. Thus, Wnt agonist 1 alleviated rat GONFH by activating β-catenin signaling, but could not rescue GONFH-like phenotype in *β-catenin^Col2ER^* mice. Nevertheless, it remains unclear how Wnt agonist 1 works to activate β-catenin signaling. Secondly, *β-catenin^OsxER^* mice presented few lipid droplet and empty lacunae but a significant decrease of bone mass in the femoral heads. Previous studies revealed that specific knockout of *β-catenin* in Osx-expressing cells promote osteoclast formation and activity^[68, 69]^, while its exact mechanism of bone mass loss still need to be verified. Thirdly, bone damage caused a poor mechanical support is the key to femoral head collapse. Although it is well known that an extensive subchondral bone destruction finally led to femoral head collapse in 6-month-old *β-catenin^Col2ER^* mice, the detailed mechanism why this subchondral bone destruction occurs with age is still unclear.

In summary, our studies revealed that β-catenin inhibition induces the imbalance in osteogenic/adipogenic differentiation of BMSCs and plays a crucial role in GONFH pathogenesis. We also provided a *Col2*-specific *β-catenin* knockout mouse model, for the first time, which mimics full spectrum of osteonecrosis phenotype of GONFH.

## Materials and methods

### Human specimen collection

Fifteen necrotic femoral head samples were collected from GONFH patients (ARCO grades III-IV) who received arthroplasty surgery. Ten non-necrotic femoral head samples were acquired from femoral neck fracture patients. All experiments with human specimens were approved by the Ethics Committee of the First Affiliated Hospital of Zhejiang Chinese Medical University (2018KL-005).

### Animals and interventions

Thirty 2-month-old male Sprague-Dawley (SD) rats were purchased from the Animal Laboratory Center of Zhejiang Chinese Medical University (Zhejiang, China, SCXK 2014-0004), and randomly divided into the sham group, the GONFH group and the treatment group (n = 10 per group). We adopted Zheng’s methods to establish a glucocorticoid-induced GONFH rat model^[9]^. Firstly, rats in the GONFH group and the treatment group received once injection of lipopolysaccharide (0.2 mg/kg body weight, Sigma, USA) by tail vein, followed by three continuous intraperitoneal injections of 100 mg/kg body methylprednisolone (MPS, Pfizer, USA). At the subsequent 6 weeks, MPS were intraperitoneally injected at the dosage of 40 mg/kg body weight three times every week. Rats in the treatment group were intravenously injected with Wnt agonist 1 (5 mg/kg body weight, Selleck, USA) at synchronous frequency with MPS.

*Col2-CreER^T2^* mice, *β-catenin^flox/flox^* mice, *Osterix-CreER^T2^*mice and *Rosa26^tdTomato^* mice were obtained from Jackson Laboratory. To map cell fate of Col2^+^ cells and Osx^+^ cells in the femoral head, *Col2-CreER^T2^;Rosa26^tdTomato^*(*Tomato^Col2ER^*) mice and *Osterix-CreER^T2^;Rosa26^tdTomato^* (*Tomato^OsxER^*) mice induced by intraperitoneally injecting with tamoxifen (TM, 1 mg/10g body weight/day, diluted in corn oil) for 5 consecutive days at the age of 2 or 4 weeks, and traced as long as 9-month-old. To further investigate the role of β-catenin in GONFH pathogenesis, *Col2-CreER^T2^;β-catenin^flox/flox^* (*β-catenin^Col2ER^*) mice and *Osterix-CreER^T2^;β-catenin^flox/flox^* (*β-catenin^OsxER^*) mice were generated and induced with 5 continuous injection of TM. After induction, Wnt agonist 1 (5 mg/kg body weight) was intravenously injected into *β-catenin^Col2ER^* mice three times every week until they were sacrificed at the age of 3 months. All animal experiments were approved by the Animal Ethics Committee of Zhejiang Chinese Medical University (No. 20190401-10).

### Stereomicroscopy observation and biomechanical testing

Femoral head tissues harvested from human patients, glucocorticoid-induced GONFH rats, *β-catenin^Col2ER^* mice and *β-catenin^OsxER^* mice were viewed under a stereomicroscope (Model C-DSD230, Nikon, Japan) to assess their appearance characteristics. The fresh femoral head tissues were placed on the test platform of a biomechanical testing machine (EnduraTec TestBenchTM system, Minnetonka, MN, USA). The axial compression load was applied at a speed of 0.5 mm/min until femoral head deformation. The loading-bearing stiffness of femoral head was calculated by testing software.

### μCT analysis

Femoral head samples were scanned with a micro-computed tomography (μCT, Skyscan 1176 Bruker μCT, Kontich, Belgium) at a resolution of 9 μm. Three-dimensional (3D) structure was reconstructed using NRecon Software v1.6 as described previously^[70]^. The region of interest in the subchondral bone was selected for morphometric analysis including bone volume fraction (BV/TV, %), average trabecular thickness (Tb.Th, mm), average trabecular number (Tb.N, 1/mm), average trabecular separation (Tb.Sp, mm) and subchondral bone defect area (%).

### Histology, immunohistochemistry and histomorphometry

Femoral head tissues were processed into 3-μm-thick paraffin section or 10-μm-thick frozen section as described previously^[71]^. After deparaffinage and rehydration, sections were stained with Alcian Blue Hematoxylin for 1 hour and then Orange G for 1 min for histological analysis. After Alcian Blue Hematoxylin (ABH) / Orange G staining, bone tissue presented yellow color and cartilage was blue. The trabecular bone area (%), fat droplet area (%) and empty lacuna rate (%) were measured using OsteoMetrics software (Decatur, GA). The frozen sections were performed with 4’,6-diamidino-2-phenylindole (DAPI) staining and hematoxylin and eosin (H&E) staining for evaluating *Cre*-recombination efficiency.

Immunohistochemistry (IHC) assay were performed on the paraffin sections according to the previously established procedures^[72]^. Briefly, sections were immersed into 0.01 M citrate buffer (Solarbio, Beijing, China) at 60L for 4 h to repair antigen. After washing with phosphate buffer (PBS) three times, the sections were incubated with primary antibodies including β-catenin (diluted 1:500, abcam, ab32572, UK), Runx2 (diluted 1:200, Abcam, ab76956, UK), alkaline phosphatase (ALP, diluted 1:200, ARIGO, ARG57422, CN), osteocalcin (OCN; diluted 1:200, Takara, UK), PPAR-γ (diluted 1:300, Abcam,ab19481, UK), CEBP/α (diluted 1:400, Huabio, EM1710-20, CA) and fatty acid-binding protein (FABP4, diluted 1:200, abcam, ab92501, UK) overnight at 4□. After incubation with secondary antibodies for 30 min at room temperature, tissue sections were stained with diaminobenzidine (DAB) solution to detect positive staining and hematoxylin for counterstaining. Mean density (IOD/Area) as the parameter of immunohistochemical quantitative analysis was evaluated using Image-Pro Plus software (Media Cybernetics, Silver Spring, USA).

### Rat BMSCs isolation and identification

BMSCs isolation were conducted as previously described^[25]^. Briefly, bone marrow cells were isolated from the proximal femur of 4-week-old SD rats. After centrifugation at 200 g for 10 min, cells were resuspended with α-MEM (Gibco, Grand Island, NY) containing 10% fetal bovine serum (FBS, Gibco, Grand Island, NY). By changing the medium 48 h later, no-adherent cells were removed and the primary BMSCs were harvested. At 100% confluence, primary cultures were detached using trypsin and subcultured at 8 × 10^3^ cells/cm^2^. A total of 50 μL third passage (P3) BMSCs suspension (4 × 10^6^/mL) were mixed with 5 μL of rat CD90-APC, rat anti-CD29-BV421, rat anti-CD45-FITC or rat CD11b-PE (BD Pharmingen, Franklin Lakes, NJ, USA) in each flow cytometry tube. After incubation at room temperature for 30 min followed by twice washes of a staining buffer, the cells were resuspend with 500 μL staining buffer for flow cytometry analysis.

### Rat BMSCs intervention and staining

To investigate the role of glucocorticoids on BMSCs differentiation, P3 BMSCs were treated with gradient concentrations of dexamethasone (Dex, 0, 10, 10^2^, 10^3^, 10^4^, 10^5^ nM) (Selleck, Shanghai, China) with or without SKL2001 (a agonist of β-catenin) (Selleck, Shanghai, China) for 72 h. For oil-red O staining, BMSCs were treated with adipogenic medium (α-MEM containing 10% FBS, 0.25 mM methylisobutylxanthine, 5 μg/mL insulin, and 50 μM indomethacin) for 72 h followed by 5 μg/mL insulin alone for an additional 48 h. After fixed in 4% paraformaldehyde for 10 min, BMSCs were stained in 60% saturated oil-red O solution for 5 min. For ALP staining, BMSCs were treated with osteogenic medium (α-MEM containing 10% FBS, 50 μg/mL ascorbic acid and 5 mM β-glycerophosphate) for 14 d, fixed in 4% paraformaldehyde for 10 min and stained with ALP staining kit (Thermo Fisher Scientific, Rockford, IL, USA) for 15 min.

### Western blot analysis

According to the appearance and CT images, two piece (about 2*2*2 cm^3^) of necrotic tissues were cut from each human necrotic femoral head, and the corresponding region from human fractured femoral head. These necrotic and non-necrotic tissues were mixed and ground into powder after processed by liquid nitrogen, respectively. Tissue powder and rat P3 BMSCs were lysed by protein extraction reagent, and the liquid supernatant containing total protein was obtained through centrifuging at 12000 g for 20 min. Then, proteins were separated by SDS-PAGE and transferred onto nitrocellulose membranes. The membranes were incubated overnight with primary antibodies including β-catenin, Runx2, Osterix, ALP, PPAR-γ and CEBP/α at 1:1000 dilution. Next, the membranes were incubated with appropriate horseradish peroxidase-conjugated secondary antibodies at 37 ℃ for 1 h. The immune complexes were detected using a chemiluminescence kit and visualized via the ChemiDoc XRS Gel documentation system (Bio-rad, CA). The gray intensity was measured by the software of Quantity One (Bio-Rad, CA).

### Statistical analysis

All data were presented as mean ± standard deviation (SD). Statistical analyses including unpaired Student’s *t*-test (two groups) and one-way ANOVA followed by Tukey’s test (multiple groups) were performed with the software of SPSS 22.0 software. \**P*<0.05, \*\**P*<0.01 was considered statistically significant.

## Data Availability Statement

Raw data are mainly attached in the current study. Any other data involving this study are available from the corresponding author upon request.

## Acknowledgements

This work has been supported by Chinese National Natural Science Foundation (grants nos. 82074469, 82104885), Natural Science Foundation of Zhejiang Province (grants nos. LQ22H270001, LY21H270008, LR23H270001), Ningbo medical key discipline (grant no. 2022-B01), Ningbo Top Medical and Health Program (grant no. 2022020102).

## Author Contributions

Di Chen, Chenjie Xia and Hongting Jin conceived this study. Chenjie Xia wrote the paper. Di Chen and Hongting Jin revised the paper. Chenjie Xia, Huihui Xu, Ping-er Wang, Liang Fang, Jiali Chen, Wenhua Yuan, Danqing Fu, Xucheng Wang performed the experiments and analyzed the data. Bangjian He, Luwei Xiao, Peijian Tong and Chengliang Wu collected the human samples.

